# Probiotic bacteria *Lacticaseibacillus rhamnosus L108* and *Lacticaseibacillus delbrueckii R2* increase lifespan and influence the expression of *Caenorhabditis elegans* longevity genes

**DOI:** 10.1101/2023.12.05.570152

**Authors:** D.S. Chelombitskaya, A.V. Teperin, S.A. Emelyantsev, E.V. Prazdnova

## Abstract

Numerous studies have shown that probiotics hold great promise in slowing down the aging process and extending life expectancy. Bacteria of the genus *Lacticaseibacillus* have been found to possess antioxidant, antimutagenic and anti-inflammatory effects, as well as the ability to regulate the expression of genes that control signaling defense mechanisms in model objects. In this study, we used the nematode *Caenorhabditis elegans* as a model organism to investigate the impact of *Lacticaseibacillus rhamnosus L108(L*.*rhamnosus L108)* and *Lacticaseibacillus delbrueckii R2* on nematode lifespan and the expression levels of genes associated with healthy aging. Our results demonstrate that the *Lacticaseibacillus rhamnosus* L108 strain exhibits antioxidant properties and increases the average lifespan of nematodes by 15%. *Lacticaseibacillus delbrueckii R2* also has a positive effect, extending the lifespan of the worms by 21.4%. Furthermore, analysis of longevity gene expression reveals a correlation between increased lifespan and activation of the insulin/insulin-like factor-1 pathway. Moreover, we observed a significant increase in the expression of the skn-1 gene, which encodes antioxidant proteins and enhances the antioxidant response. Our findings suggest that the expression of the skn-1 gene and the transcription factor SKN-1 are associated with activation of the p38 MAPK signaling pathway. Therefore, it seems that probiotic bacteria *Lacticaseibacillus rhamnosus L108 and Lactobacillus delbrueckii R2* have a positive effect on lifespan due to increased expression of genes that underlie the regulation of conserved signaling pathways related to host defense.

## Introduction

Numerous studies have been dedicated to discovering products that can slow down the aging process in the body. The aging of our bodies is a natural physiological phenomenon that leads to a decline in the functioning of all organ systems and the occurrence of changes at the tissue and cellular levels. These changes include increased genomic instability, telomere attrition, epigenetic alterations, disruptions in protein metabolism, impaired nutrient sensitivity, mitochondrial dysfunction, and overall cellular aging.

As a result, the investigation of probiotic microorganism groups has emerged as a promising area of research. Back in the early 20th century, I.I. Mechnikov proposed the idea that using live lactic acid bacteria as a dietary supplement could enhance immunity and extend human lifespan.

Today, probiotics are known to have positive effects on host health through various molecular mechanisms. They possess antioxidant, antimicrobial, and anti-inflammatory properties. Exploring the ability of probiotic bacteria from the Lactobacillus genus in regulating the expression of genes that control protective signaling mechanisms, which contribute to lifespan extension and antioxidant response, requires the use of simple model systems. The nematode *Caenorhabditis elegans* is an ideal model organism for studying the molecular mechanisms of antioxidant defense and their genetic contributions due to its ease of cultivation in vitro, transparent skin, short life cycle, high productivity, and traceable genetics. Moreover, it was discovered that signaling mechanisms for protection and longevity genes in *Caenorhabditis elegans* share around 60-80% homology with humans.

The utilization of probiotics to counter premature aging is a promising field of scientific inquiry. Research has shown that probiotics promote the production of antimicrobial substances, enhance the integrity of the host’s mucosa, and prevent the proliferation of harmful microorganisms. In this study, we assessed the probiotic strains *Lacticaseibacillus rhamnosus L108 and Lacticaseibacillus delbrueckii R2*, which were isolated from traditional dairy products, to determine their impact on the lifespan of nematodes and the relationship between molecular defense mechanisms and intestinal microflora.

Experimental evidence confirms that probiotic bacteria from the *Lactobacillus* and *Bifidobacterium* genera have antioxidant effects and can synthesize antioxidant complexes to combat oxidative stress. When consumed as food, probiotics can extend lifespan and mitigate oxidative stress through the DAF-2/DAF-16 insulin/insulin-like factor-1 (IIS) signaling pathway. This signaling system is involved in regulating lipid and glucose metabolism, protein synthesis, and cell proliferation.

The objective of our study was to evaluate the influence of the probiotic bacterium *Lacticaseibacillus rhamnosus L108* on the lifespan of *Caenorhabditis elegans* and the expression of longevity genes associated with the molecular pathways of insulin/insulin-like growth factor-1 (IIS) and p38 mitogen-activated protein kinase (p38 MAPK).

## Materials and methods

Probiotic strains. The bacteria were obtained from the collection of the laboratory of experimental mutagenesis of the Academy of Biology and Biotechnology named after. D.I. Ivanovsky Southern Federal University. Their antioxidant and antimutagenic properties are characterized in the work. [21]

### 1. Cultivation of *Lacticaseibacillus rhamnosus L108, Lacticaseibacillus delbrueckii R2*

Strains *Lacticaseibacillus rhamnosus L108 and Lacticaseibacillus delbrueckii R2* were cultured in liquid MRS medium for 48 hours.

Composition of MRS medium per 1 liter in grams: proteose peptone - 10.0; meat extract - 10.0; yeast extract - 5.0; glucose - 20.0; polysorbate 80 - 1.0; ammonium citrate - 2.0; sodium acetate - 5.0; magnesium sulfate - 0.1; manganese sulfate - 0.05; disubstituted potassium phosphate - 2.0. pH (at 25°C) 6.5, 1 g of Tween 80. pH of the medium is within 6.2 (+/- 0.2) [9]. Cultivation was carried out under anaerobic conditions in a thermostat at a temperature of 37□ C.

### 2. Method for cultivating the nematode Caenorhabditis elegans

Nematodes *Caenorhabditis elegans* were cultured on plastic Petri dishes with a diameter of 90 mm on NGM agarose medium, inoculated with a lawn of E. coli OP50, in a thermostat at a temperature of 25□□ C. The E. coli OP50 strain was cultivated in liquid LB medium in a thermostat at a temperature of 37□ C for 24 hours.

To obtain a population of the same age, C. elegans nematodes were subjected to chemical synchronization. Worms were collected from NGM agarose medium by rinsing using M9 buffer into sterile 1.5 ml plastic Eppendorf tubes. Eppendorfs were centrifuged at 4000 g for 60 seconds, then the supernatant was carefully removed without disturbing the pellet. 1 ml of buffer M9 was added and this procedure was repeated 3 times to purify the nematodes from bacterial cells. Eggs were extracted in a solution containing 0.5 M NaOH with 1% Na-hypochlorite and then washed with buffer M9 at least three times. Eggs were transferred to inoculated Petri dishes with NGM E. coli OP50 and incubated overnight at 20°C to obtain synchronized L1 larval stage animals.

### 3. Analysis of the lifespan of the nematode *Caenorhabditis elegans*

For the first screen, adults were selected from a homogeneous population and transferred to lawn plates with *E. coli OP50, L. rhamnosus L108 and L. delbrueckii R2*. The nematodes on the standard lawn, E. coli OP50, were considered the control group, and the worms on the lawn, *L. rhamnosus L108 and L. delbrueckii R2*, were considered the experimental groups.

For lifespan analysis, nematode eggs obtained by chemical synchronization of *C. elegans* with a solution of 0.5 M NaOH and 1% Na-hypochlorite were used. The eggs were incubated in M9 buffer in an incubator at 25 C for 24 hours and then dropped onto NGM agarose plates pre-coated with the probiotics *Lacticaseibacillus rhamnosus L108, Lacticaseibacillus delbrueckii R2 and E. coli OP50*. Nematodes in the lawn with E. coli OP50 were considered as a control. Within 2-3 days, the worms were transferred to new plates to distinguish the offspring from the studied individuals. The total time to analyze the lifespan of *Caenorhabditis elegans* without oxidative stress on probiotic lawn was 21 days. Nematodes were checked for signs of vital activity using a platinum wire held close to the body. Worms that did not respond to the stimulus were considered dead.

### 4. RNA extraction and quantitative real-time PCR (qRT-PCR)

Total RNA of nematodes was extracted using commercial kits for RNA isolation from whole blood, cell cultures, and samples of animal and plant tissues “RNA-EXTRAN” from SINTHOL and “ExtraRNA” from Evrogen. For additional purification of the resulting RNA, a CleanRNA Standard kit with spin columns was used. Reverse transcription of the resulting RNA samples was carried out using the “MMLV RT kit” from EuroGen.

The synthesized complementary DNA was subjected to qRT-PCR analysis using qPCRmix-HS SYBR from EuroGen in an ANK-32-M qRT-PCR amplifier. Oligonucleotides for RT-PCR were designed using the Primer3 tool. The Ct values obtained during the expression of the studied genes were compared with respect to the reference gene gsr-1. Gene expression measurements are obtained by calculating relative expression levels using the ΔΔCt method.

## Results

Lifespan analysis of the nematode *Caenorhabditis elegans*

Before the experiment, it was found that nematodes exhibit normal growth and reproduction when grown both on probiotic strains and on E. coli OP50.

The probiotic bacterium *L. rhamnosus L108* has been shown to increase the lifespan of *C. elegans* nematode by 9.6% (p<0.05) and average lifespan by 15% (p<0.05) compared to a control group of worms, that fed on the lawn E. coli OP50 (Fig. 1). In the same model, *L. delbrueckii R2* increases nematode lifespan by 21.4% (p<0.05) and average lifespan by 39.1% (p<0.05) compared to the control group (Fig. 2).

**Figure 1.**
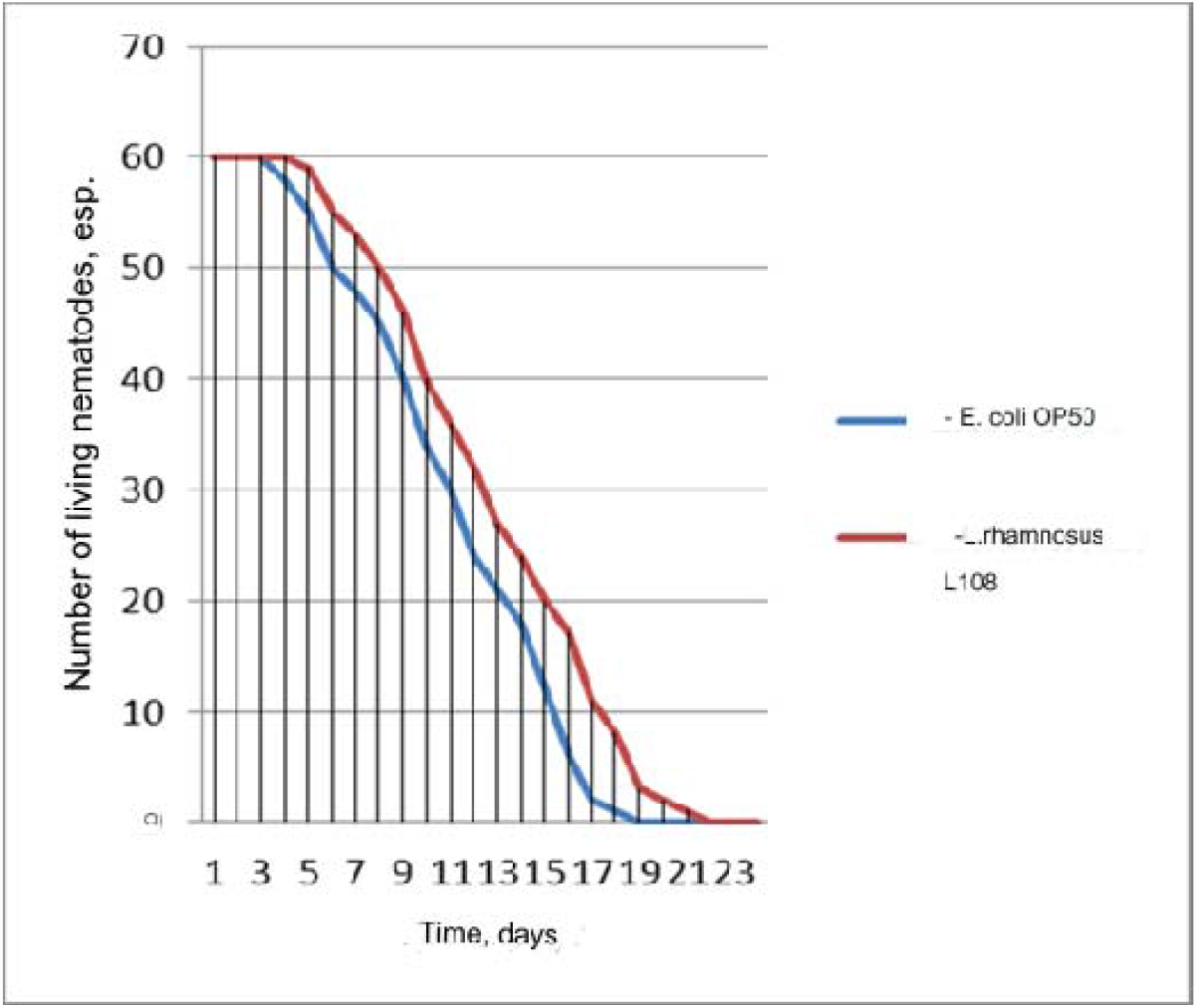
Survival curves of *C. elegans* nematodes cultured on control plates with *E. coli OP50* lawn and the studied probiotic *L. rhamnosus L108*

**Figure 2.**
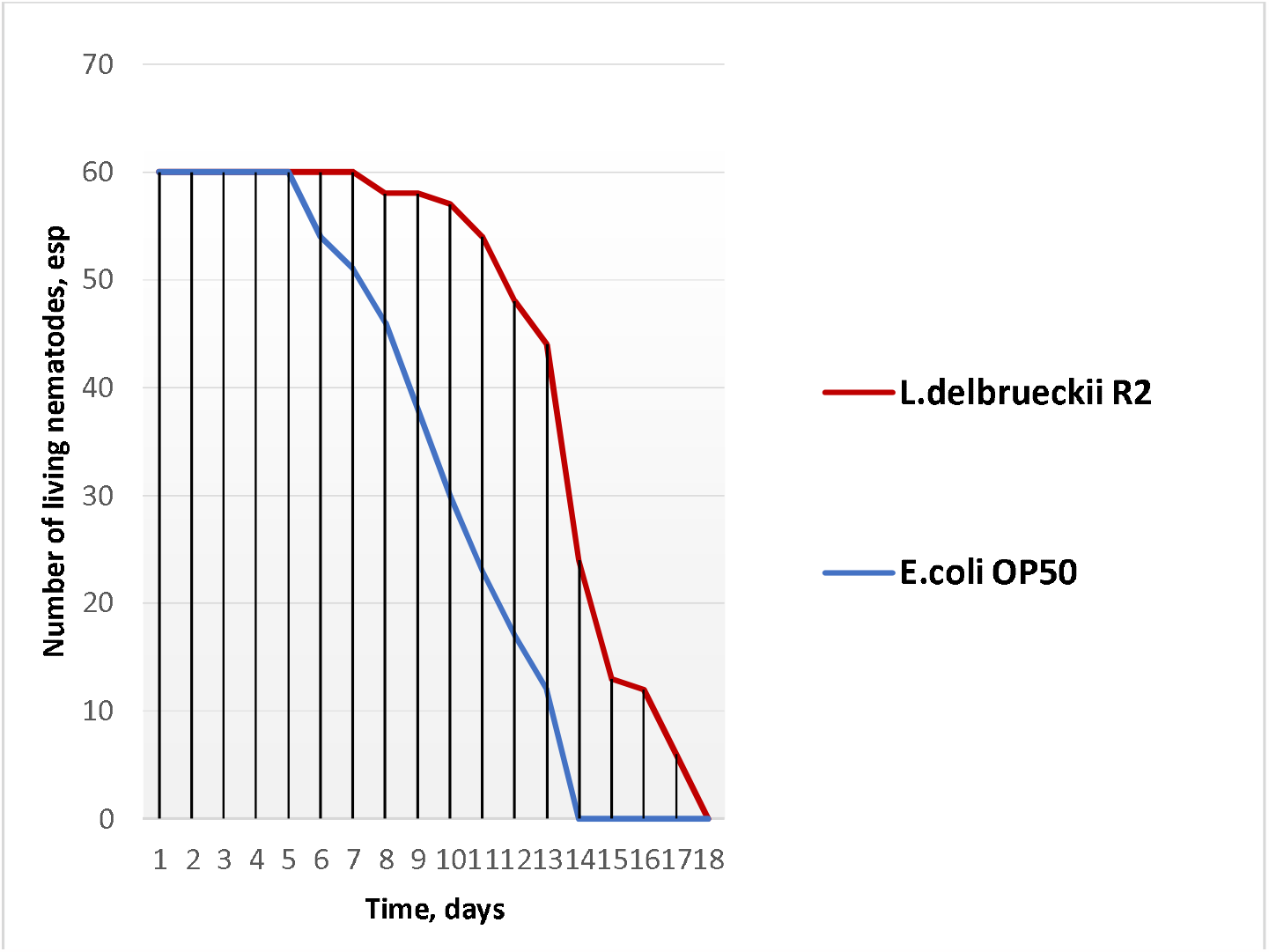
Survival curves of *C. elegans* nematodes cultured on control plates with *E. coli OP50* lawn and the studied probiotic *L. delbrueckii R2*

To identify the effect of the probiotic L. rhamnosus L108 on the genetic apparatus of the nematodes *C. elegans*, we examined the expression of the sir-2.1, skn-1, daf-16 genes, which underlie the regulation of the conservative signaling protective pathways insulin/insulin-like factor-1 and p38 MAPK.

Gene expression analysis was carried out at three time points: after the 1st, 3rd and 7th day of feeding *Lactocasiebacillus rhamnosus L108*. Figure 3 shows an increase in the expression level of the sir-2.1 gene after daily feeding with a probiotic by 9%, after 3 days - by 34%. After the 7th day, a decrease in expression by 43.6% (p < 0.05) is observed.

**Fig. 3.**
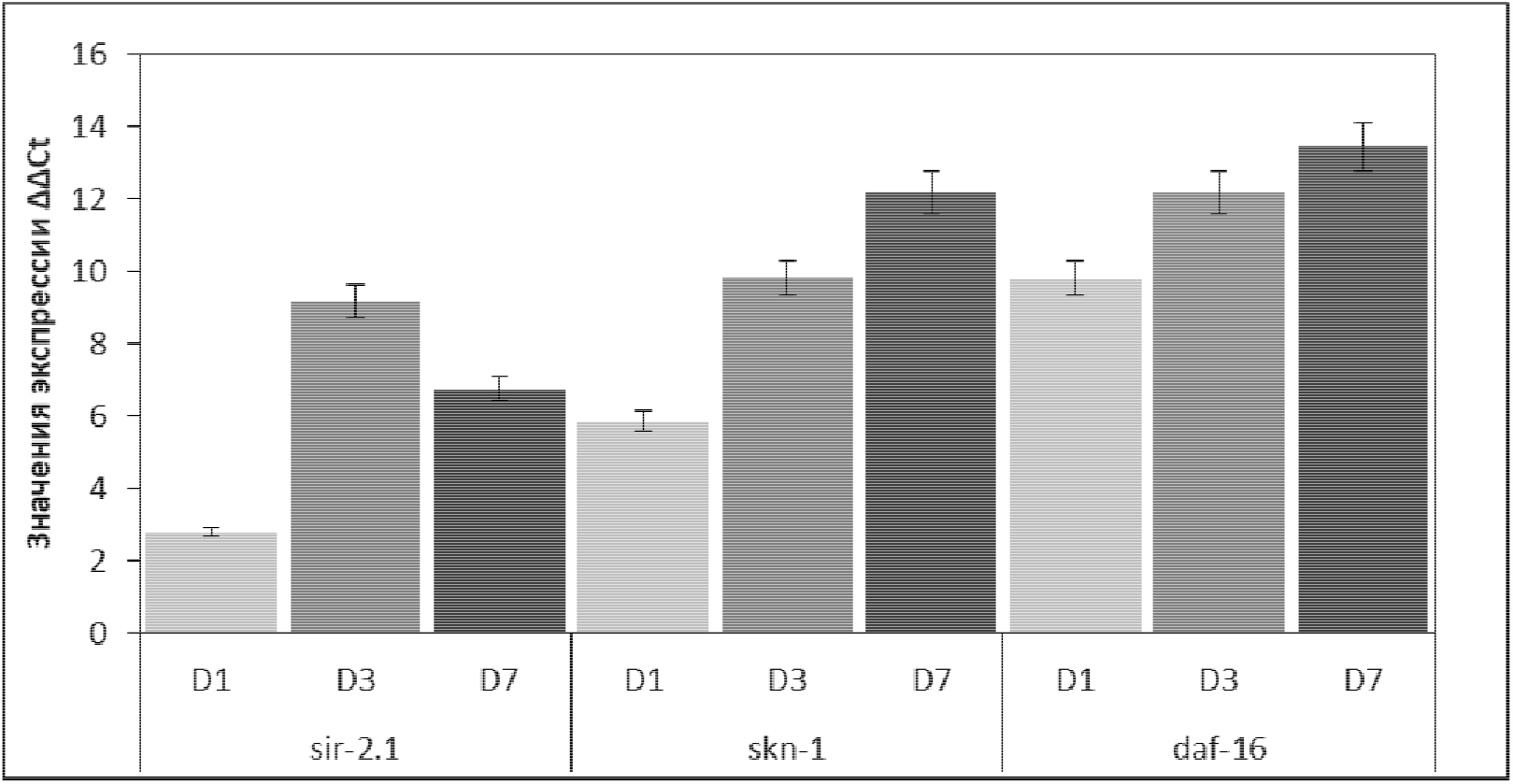
Histogram based on the relative expression of the sir-2.1, skn-1 and daf-16 genes after feeding the nematodes *C. elegans* with the probiotic *Lactocasiebacillus rhamnosus L108*. Note: the expression levels of the studied *C. elegans* genes are shown after the 1st (D1), 3rd (D3) and 7th (D7) days of consumption of *L. rhamnosus L108*.

**Figure 4.**
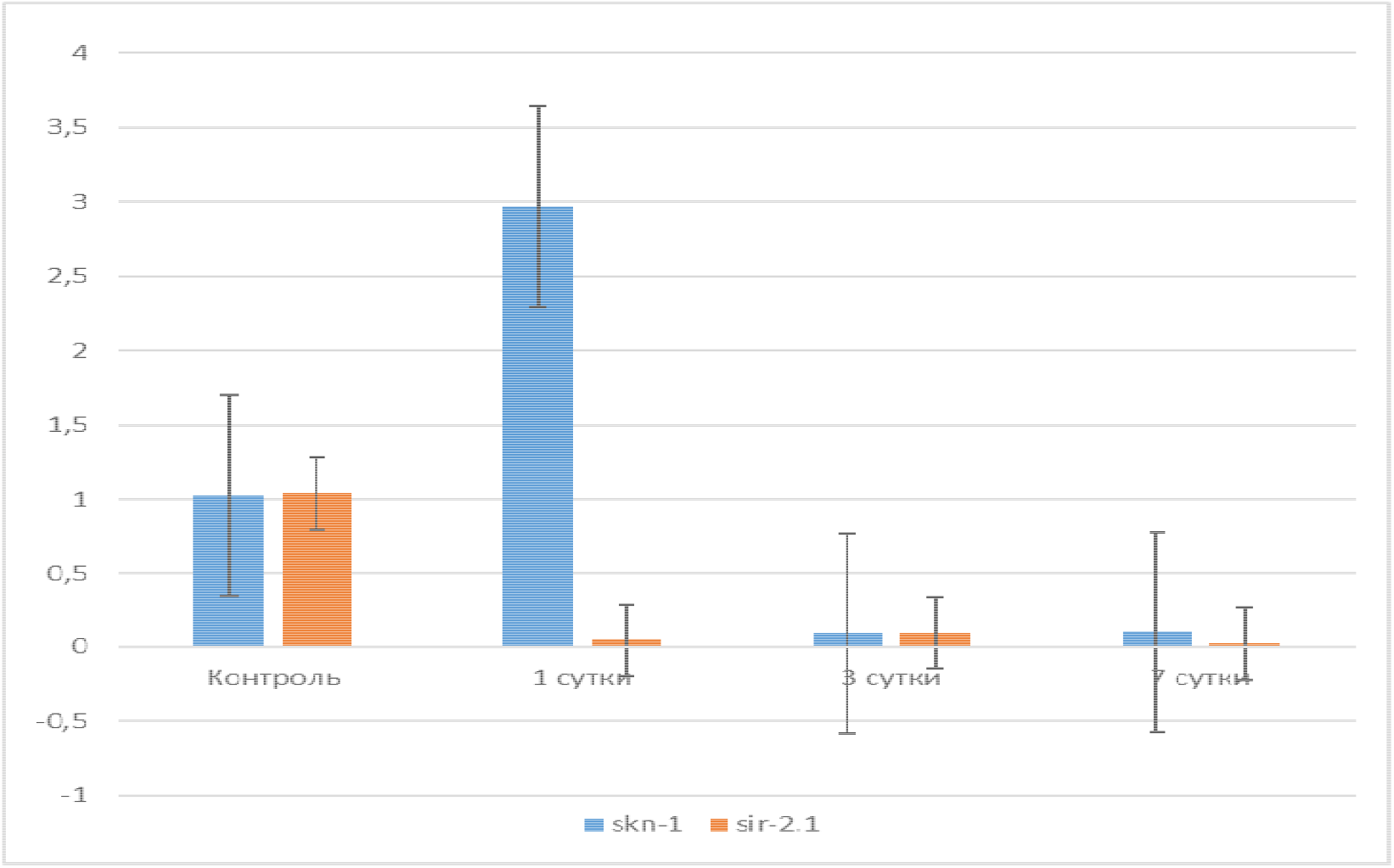
Bar graph according to the relative expression of the sir-2.1 and skn-1 genes on days 1, 3 and 7 when *C. elegans* nematodes were fed the control *E. coli strain OP50* and the test strain *L. delbrueckii R2*

Expression of the skn-1 gene increased exponentially at all three time points by 43%, 109% and 184% (p < 0.05). Analysis of daf-16 gene expression data showed an increase in the expression level by 0.2% after the 1st day of feeding with a probiotic, after 3 days - by 7.2%, after 7 days - by 8.5% (p< 0.05).

In addition, we investigated which signaling pathway is affected by the introduction of the probiotic *L. delbrueckii R2* into the diet of the nematode *Caenorhabditis elegans*. Graph 2 shows a 3-fold increase in skn-1 gene expression compared to the control group of worms. At the same time, a significant decrease in its expression was observed at the other two control points. Expression of the sir-2.1 gene in the experimental group was also significantly reduced at all three time points compared to the control group of nematodes.

## Discussion

The use of probiotics to combat premature aging and their potential antioxidant properties is an exciting area of scientific research [16]. Various studies have examined the effects of probiotics on the lifespan of *C. elegans*, a model organism commonly used in aging research [16,17]. In one study, *C. elegans* were fed *L. rhamnosus CNCM I-3690* for 21 days and compared to worms fed *E. coli OP50*. The results indicated that feeding the CNCM I-3690 strain increased the average lifespan of the worms by 3 days compared to those fed OP50. The protective effect of *L. rhamnosus* became apparent after 7 days of feeding, and a 20% increase in worm survival was observed after 15 days [19].

Another study focused on *L. delbrueckii R2* and its effects on the transcription factor SKN-1, which plays a role in enhancing the transcriptional activity of various antioxidant genes: sod-1, gcs-1, clk-1, trx-1, hsp70, detoxification genes and xenobiotics [18]. The study found that L. delbrueckii R2 stimulated SKN-1 activity and subsequently increased the expression of antioxidant genes involved in maintaining cellular homeostasis [18,19].

Additionally, the probiotics *L. rhamnosus L108 and L. delbrueckii R2* were found to increase the lifespan of C. elegans and regulate the aging process through insulin/insulin-like growth factor-1 (IIS) and p38 MAPK signaling pathways [20].

Aging is a complex physiological process characterized by disruption of the functioning of all molecular mechanisms within the body [22]. Its main features are: increased genomic instability, telomere attrition, epigenetic changes, impaired protein metabolism, impaired nutrient sensitivity, mitochondrial dysfunction and general cellular aging [22,23].

Aging is closely related to oxidative stress [24], which is a promising, although not the only, direction in the search for geroprotectors [24]. Overall, the findings suggest that probiotics with antioxidant properties have the potential to be effective geroprotectors in vivo [24]. Further research is needed to unravel the specific mechanisms underlying these effects and to explore the potential therapeutic applications of probiotics in combating premature aging [24].

In this study, among the three genes studied, the highest level of expression was recorded for the skn-1 gene, which underlies the regulation of the p38 MAPK signaling pathway. The p38 MAPK signaling pathway is known to increase the activity of the immune response of worms by activating a transcription factor known as SKN-1, encoded by the skn-1 gene, as well as antioxidant and detoxification genes (sod-1, gcs-1, clk-1, trx -1, hsp70 and hsp16.2) [26,27]. A smaller effect is observed when analyzing the expression of the sir-2.1 and daf-16 genes, which underlie the regulation of the DAF-2/DAF-16 insulin/insulin-like factor-1 (IIS) signaling pathway. This signaling system is involved in the regulation of lipid metabolism, glucose metabolism, protein synthesis and cell proliferation [28]. The mechanisms by which these strains directly affect lipid homeostasis in worms remain unknown [28]. Despite this, a possible correlation between these results and the increased lifespan of worms fed *L. rhamnosus L108 and L. delbrueckii R2* cannot be ruled out [29,30]. It is conceivable that some bacterial ligands may trigger the transcription factor DAF-16 by dephosphorylation or another mechanism that allows DAF-16 to translocate into the nucleus, but this requires further confirmation in vivo [31].

## Conclusion

Thus, in our study, we determined the positive effect of *L. rhamnosus L108 and L. delbrueckii R2* on the lifespan of *Caenorhabditis elegans* and the expression of the longevity genes sir-2.1 and skn-1, which underlie the regulation of conserved protective signaling pathways insulin/insulin-like factor-1 and p38 MARK. The highest level of expression is observed in the skn-1 gene, which is part of the p38 MAPK defense mechanism.

Thus, during the experiments, the strain *Lacticaseibacillus rhamnosus L108*, which had an antioxidant effect and increased the average lifespan of the nematode *Caenorhabditis elegans* by 15% and significantly increased the expression of the skn-1 p38 MAPK signaling pathway gene. *Lacticaseibacillus delbrueckii R2* also had a positive effect and increased the lifespan of the worms by 21.4%.

## Acknowledgments

The research was financially supported by the Ministry of Science and Higher Education of the Russian Federation (№ FENW-2023-0008). The study was carried out with the support of the Laboratory of Experimental Mutagenesis of the Academy of Biology and Biotechnology named after. D.I. Ivanovsky Southern Federal University.

## Availability of Data and materials

All data generated or analyzed during this study are included in this published article and comply with research standards.

